# Reduced plasticity in coupling strength in the SCN clock in aging as revealed by Kuramoto modelling

**DOI:** 10.1101/2021.09.13.460004

**Authors:** Anouk W. van Beurden, Janusz M. Meylahn, Stefan Achterhof, Johanna H. Meijer, Jos H. T. Rohling

## Abstract

The mammalian circadian clock is located in the suprachiasmatic nucleus (SCN) and consist of a network of coupled neurons, which are entrained to the environmental light-dark cycle. The phase coherence of the neurons is plastic and driven by the length of the day. With aging the capacity to behaviorally adapt to changes in the light regime reduces. The mechanisms underlying photoperiodic adaptation are largely unknown, but are important to unravel for the development of novel interventions to improve the quality of life of the elderly. We analyzed the neuronal synchronization of PER2::LUC protein expression in the SCN of young and old mice entrained to either long or short photoperiod and used the synchronization levels as input for a two-community noisy Kuramoto model. With the Kuramoto model we estimated the coupling strength between and within neuronal subpopulations. The model revealed that the coupling strength between and within subpopulations contributes to photoperiod induced changes in the phase relationship among neurons. We found that the SCN of young mice adapts in coupling strength over a large range, with low coupling strength in long photoperiod and higher coupling strength in short photoperiod. In aged mice we also found low coupling strength in long photoperiod, but strongly reduced capacity to reach high coupling strength in short photoperiod. The inability to respond with an increase in coupling strength shows that manipulation of photoperiod is not a suitable strategy to enhance clock function with aging. We conclude that the inability of aged mice to reach high coupling strength makes aged mice less capable to seasonal adaptation than young mice.

**Author Summary:** Circadian clocks drive daily rhythms in physiology and behavior. In mammals the clock resides in the suprachiasmatic nucleus (SCN) of the hypothalamus. The SCN consist of a network of coupled neurons which are synchronized to produce a coherent rhythm. Due to plasticity of the network, seasonal adaptation to short winter days and long summer days occurs. Disturbances in circadian rhythmicity of the elderly have negative health effects, such as neurodegenerative diseases. With the rise in life expectancy this is becoming a major issue. In our paper, we used a model to compare the neuronal coupling in the SCN between young and old animals. We investigated whether exposure to short photoperiod can strengthen coupling among clock cells, and thereby clock function, in old animals. We observed that this is not possible, indicating that simple environmental manipulations are not an option. We suggest that receptor targeted interventions are required, setting the path for further investigation.

## Introduction

Many organisms increase their chance of survival and reproduction by anticipating seasonal changes in temperature and food availability. Internal clocks drive the circadian and seasonal rhythms, responsible for physiological and behavioral adaptation. In mammals, the endogenous clock is located in the suprachiasmatic nucleus (SCN) of the anterior hypothalamus in the brain. The SCN is a relatively small structure that consist of approximately 20.000 neurons [1]. Generation of circadian rhythms occurs autonomously in all individual neurons and is based on a negative feedback loop between clock genes and their protein products [2-4]. This renders a population of autonomous oscillators that have to synchronize in order to produce a coherent rhythm at the population level [5,6]. The phase coherence is plastic, and programmed by the length of the day, allowing the animal to adapt to the seasonal cycles [7-10].

How phase coherence is established at the network level is relevant for seasonal adaptation and breeding, but also for understanding clock disturbance in aging [11]. Although it is known that differences in the phase relationship between neurons underlie photoperiodic adaptation, the mechanism is unknown. One may intuitively expect that a decrease in coupling strength leads to a broadened phase distribution. Alternatively, phase differences can be driven by an active process, for example due to repulsive coupling between subpopulations of SCN neurons [12]. In such a scenario, coupling within these subpopulations could be equally strong in long and short photoperiod.

Subpopulations of SCN neurons form phase clusters, that map approximately to the core and shell SCN, and to the anterior and posterior SCN [7,13,14]. The question addressed in this study is whether we can explain the changes in synchronization of the activity phase of the neurons between different photoperiods by changes in coupling strength. Particularly we will investigate the ability of older mice to adjust to changes in daylength.

For optimal functioning of the SCN a combination of molecular (e.g. clock gene expression), cellular (e.g. electrical activity) and network (e.g. neurotransmitters) elements are important [15]. Neurotransmitters play a crucial role in synchronizing the neurons in the SCN. An age-related decline in expression of neurotransmitters has been reported [16], probably causing reduced communication among neurons in the aged SCN [17]. It has been suggested that weakened circadian rhythmicity of the elderly have negative health effects, and is causal to a broad array of diseases [18]. Therefore, strengthening the clock in the aged is important, and strategies to do so rely on an identification of underlying mechanisms. Here we investigated whether mechanisms underlying age-related changes in synchronization are the same as mechanisms underlying photoperiod induced changes in synchronization.

We used data from bioluminescence imaging of single-cell PER2::LUC gene expression rhythms as input for a Kuramoto model [19,20] to estimate the coupling strength within and between neuronal subpopulations in young and old mice entrained to long (LP, LD 16:8) and short (SP, LD 8:16) photoperiod [7,15]. Neuronal subpopulations of the SCN were identified with an unbiased clustering algorithm [21]. We took into account that the coupling strengths are not the same within and between the different neuronal subpopulations, since it is known that in the SCN the core projects densely to the shell while the shell projects only sparsely to the core [4]. We found evidence that coupling strength within and between subpopulations contributes to photoperiod induced changes in the phase relationship among neurons. Moreover, the SCN of young animals is able to adjust the coupling strength over a larger range, with lower coupling strength in long photoperiod and higher coupling strength in short photoperiod. Old animals appear to have a diminished range in coupling strength, and particularly are unable to increase coupling in short photoperiod.

## Results

### Synchronization of PER2::LUC rhythms in the SCN

We calculated the order parameter (*r)* and peak time dispersion from the smoothed bioluminescence traces (Fig 1A) for all SCN slices in the different experimental conditions. To test whether the order parameter is an appropriate measure for synchronization we calculated the Pearson correlation coefficient between *r* and peak time dispersion, which was taken as a measure for synchronization in [7,15]. The correlation coefficient showed a strong negative correlation between *r* and peak time dispersion (R=-0.91; Fig 1B), which is expected as high dispersion should lead to lower synchrony (*r*). Furthermore, we compared the values of *r* between the different experimental conditions. Independent t-tests showed that the *r* value was always significantly higher in SP than in LP in both young and old mice (young anterior, LP: 0.49±0.23, n=4, young anterior, SP: 0.87±0.04, n=5, p<0.05; young posterior, LP: 0.77±0.12, n=4, young posterior, SP: 0.91±0.03, n=5, p<0.05; old anterior, LP: 0.53±0.23, n=7, old anterior, SP: 0.80±0.08, n=10, p<0.01; old posterior, LP: 0.77±0.06, n=9, old posterior, SP: 0.83±0.04, n=10, p<0.05; Fig 1C). These results are in agreement with the results of [15].

**Fig 1.**
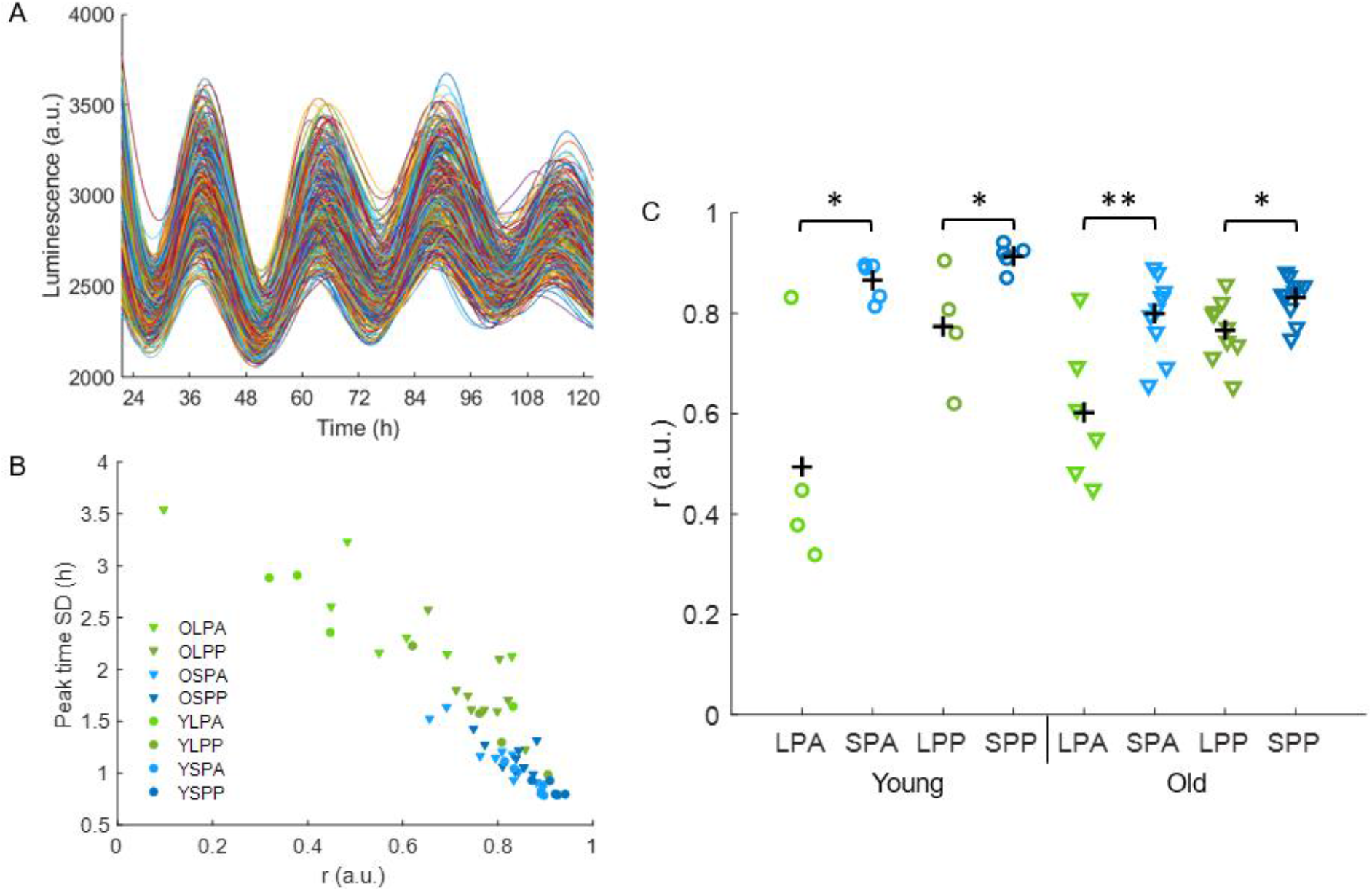
Synchronization of the SCN. (A) Example of smoothed intensity traces of PER2::LUC expression from single-cells in the anterior SCN of a young mice in short photoperiod. (B) Pearson correlation between r and peak time dispersion for all recordings (n=54, R=-0.91). (C) The order parameter r is calculated for all slices and is shown for anterior and posterior slices in long (green dots) and short photoperiod (blue dots) in young and old mice. The black crosses indicate the mean; *p<0.05, **p<0.01.

### Coupling strength and noise estimation

To determine the coupling strength (*K*) between the neurons in the SCN for the different experimental conditions, we used *r* as input to the one-community Kuramoto model. Furthermore, we estimated the level of noise (*D*) in the model. For both the coupling strength and the noise we calculated for each slice an upper and lower bound (Fig S1). A one-sample Kolmogorov-Smirnov test showed that *K* and *D* were not normally distributed (p>0.05). To compare the bounds of *K* and *D* between the experimental conditions we used non-parametric independent-samples median tests. The lower and upper bound of *K* is always significantly higher in SP than LP (p<0.05), except for the upper bound of the posterior SCN in old mice (Fig S1A). There were no significant differences in the lower and upper bound of *D* between the experimental conditions (Fig S1B). Next, the ranges between the medians of the upper and lower bounds for *K* and *D* in the different experimental conditions were calculated (Fig 2). For young mice the range for *K* in LP does not overlap with the range for *K* in SP. Therefore the coupling strength is definitely higher in SP than LP in young mice. For old mice the range of *K* in SP lies within the upper half of the range of *K* in LP, which indicates that *K* again is higher in SP than LP, although this is not significant. The range between the upper and lower bound for *D* is larger for LP than SP in both young and old mice, however the range does not differ significantly between the experimental conditions. The mean value between the upper bound and lower bound of *D* is close to one for all experimental conditions. This shows that *D* will not significantly impact the results of the two-community Kuramoto model, provided that *D* has a constant value that is independent of the synchronization level.

**Fig 2.**
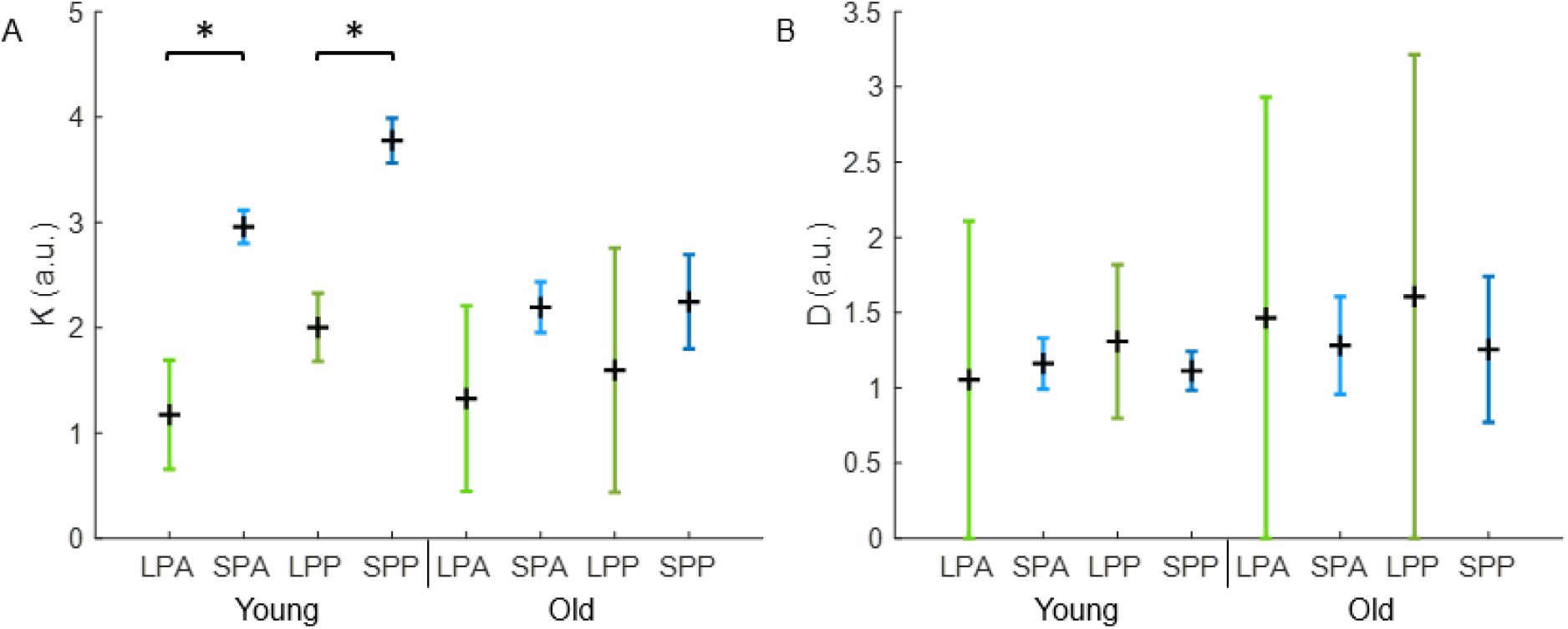
Range of *K* and *D* in different experimental conditions. (A) Range for the coupling strength between neurons in anterior and posterior slices in long (green) and short (blue) photoperiod in young and old mice. The range is based on the distance between the median of the upper and lower bound of *K* in each condition. The black cross indicates the mean of the range. (B) Range for the noise term in anterior and posterior slices in long (green) and short (blue) photoperiod in young and old mice. The range is based on the distance between the median of the upper and lower bound of *D* in each condition. The black cross indicates the mean of the range.

### Synchronization of the neuronal subpopulations

Next, we calculated the order parameter for the two neuronal subpopulations, that were identified using an unbiased community detection algorithm [21]. Note that the spatial distribution of the neuronal subpopulations only partially corresponds with the division of the SCN in dorsomedial (shell) and ventrolateral (core) SCN based on neuropeptide content [24] and differs between the anterior and posterior slices (Fig 3A). From now on we will refer to the ventromedial cluster from anterior slices and the medial cluster from posterior slices the *medially oriented cluster*. We will refer to the dorsolateral cluster from anterior slices and the lateral cluster from posterior slices the *laterally oriented cluster* for simplicity. Paired-sampled t-tests showed that *r* was always significantly higher in each of the neuronal subpopulations than in the SCN as a whole (p<0.05, result not shown). For the medially oriented cluster there was only a significant difference in *r* between LP and SP in the anterior SCN of young mice (young anterior, LP: 0.66±0.12, n=4, young anterior, SP: 0.92±0.02, n=5, p<0.05; Fig 3B). For the laterally oriented cluster *r* was significantly higher in SP than in LP in nearly all conditions, except for the posterior SCN of young mice (young anterior, LP: 0.78±0.08, n=4, young anterior, SP: 0.95±0.01, n=5, p<0.01; young posterior, LP: 0.85±0.09, n=4, young posterior, SP: 0.92±0.02, n=5, p=0.286; old anterior, LP: 0.74±0.08, n=7, old anterior, SP: 0.92±0.03, n=10, p<0.01; old posterior, LP: 0.80±0.08, n=9, old posterior, SP: 0.89±0.04, n=10, p<0.01; Fig 3C).

**Fig 3.**
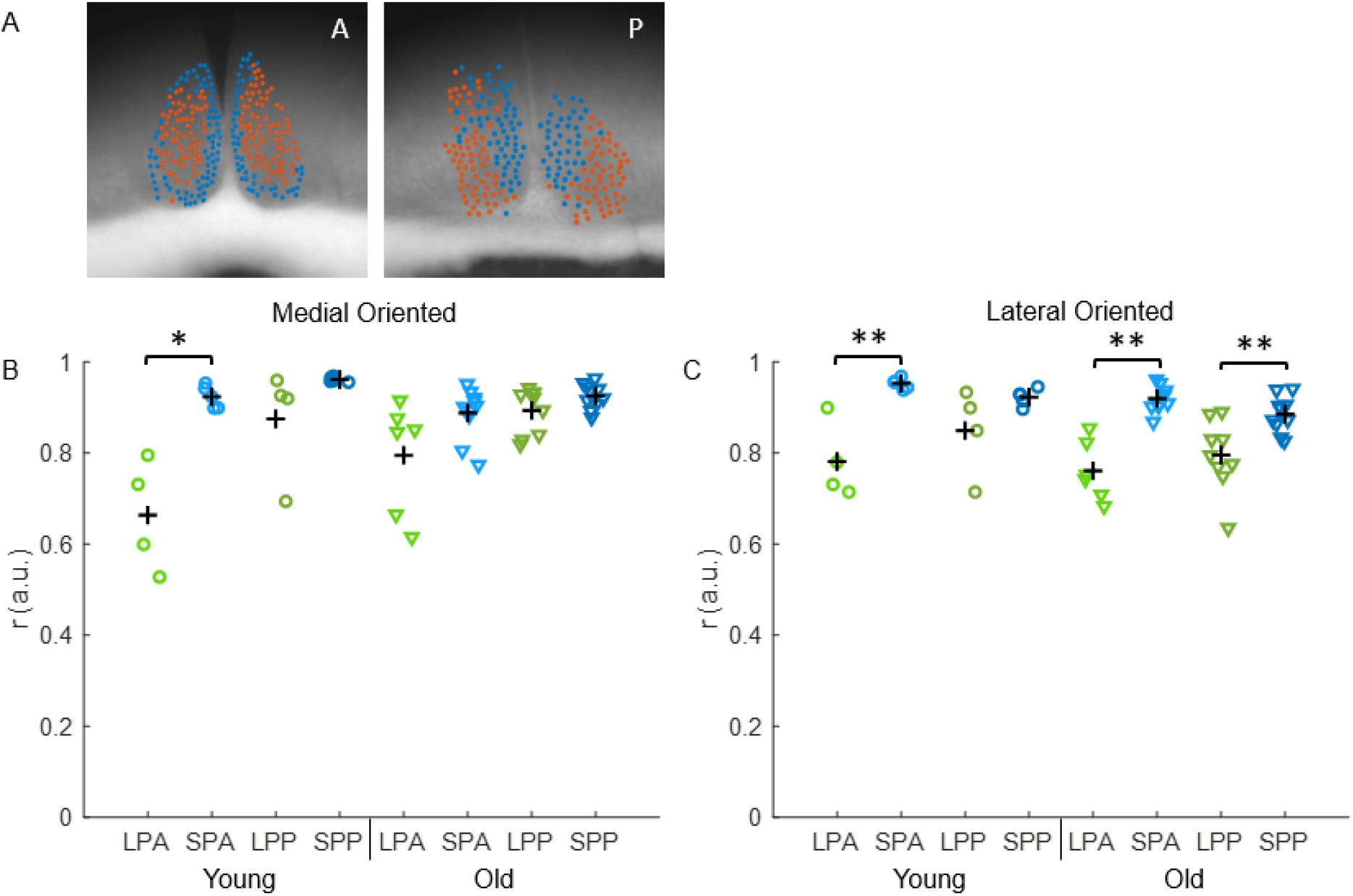
Synchronization in the SCN neuronal subpopulations. (A) Cell location projected on bright field image of the anterior (left) and posterior (right) SCN. The blue cells represent the medial oriented cluster and the orange cells the lateral oriented cluster. (B/C) The order parameter is calculated for both subpopulations of all slices and is shown for anterior and posterior slices in long (green dots) and short photoperiod (blue dots) in young and old mice. The black crosses indicate the mean of the experimental condition. Fig B shows the result of the medial oriented cluster and Fig C of the lateral oriented cluster; *p<0.05, **p<0.01.

### Estimation within and between community coupling strength

Next, we used the order parameter as calculated for the subpopulations in the different experimental conditions as input for the extended Kuramoto model [19,20]. We made the assumption that *D=1* for all experimental conditions, since the changes in *D* were minor in the results of the one-community Kuramoto model. To estimate the relationship between *K*_*1*_ and *L*_*1*_ and *K*_*2*_ and *L*_*2*_ the following equations, which are based on the extended Kuramoto model [19,20] were solved for each experimental condition. The equation

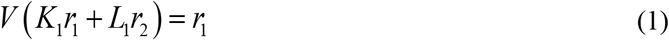

shows the relation between *K*_*1*_ and *L*_*1*_. Where *K*_*1*_ represents the coupling strength within the medially oriented cluster, *L*_*1*_ represents the interaction strength from the lateral oriented cluster to the medial oriented cluster and *r*_*1*_ is the order parameter for the medial oriented cluster. And the equation

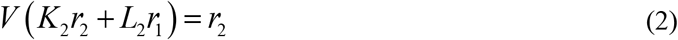

shows the relation between *K*_*2*_ and *L*_*2*_. Where *K*_*2*_ represents the coupling strength within the lateral oriented cluster, *L*_*2*_ represents the interaction strength from the medial oriented cluster to the lateral oriented cluster and *r*_*2*_ is the order parameter for the lateral oriented cluster. Fig 4 shows a simplified representation of the model.

**Fig 4.**
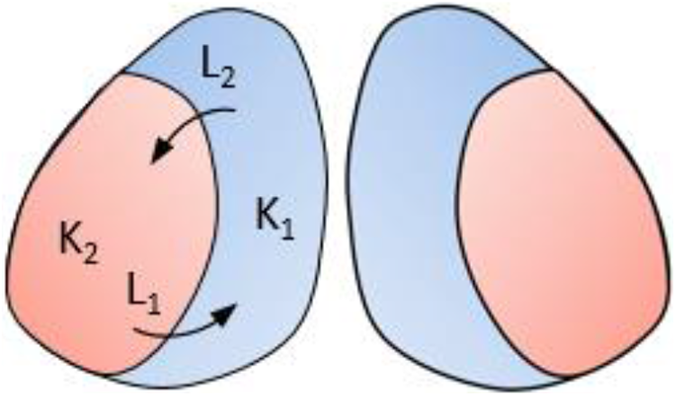
Simplified representation of the two-community Kuramoto model. The blue area represent the medial oriented cluster in which the coupling strength is denoted by *K*_*1*_ and the orange area represents the lateral oriented cluster in which the coupling strength is denoted by *K*_*2*_. *L*_*1*_ shows the interaction strength from the lateral oriented cluster to the medial oriented cluster and *L*_*2*_ shows the interaction strength from the medial oriented cluster to the lateral oriented cluster.

In Fig 5 the relationship between *K* and *L* is shown for the different experimental conditions. For both subpopulations we found a negative linear relation between *K* and *L*. The coupling strength (*K*) within a neuronal subpopulation is always positive and the interaction strength (*L*) between the neuronal subpopulations can both be positive or negative, in which a negative strength indicates an inhibitory connection. A high level of synchronization within a cluster can either be reached with high coupling strength within the cluster and when the cluster receives low interaction strength from the other cluster or with moderate coupling strength within the cluster and moderate interaction strength from the other cluster.

**Fig 5.**
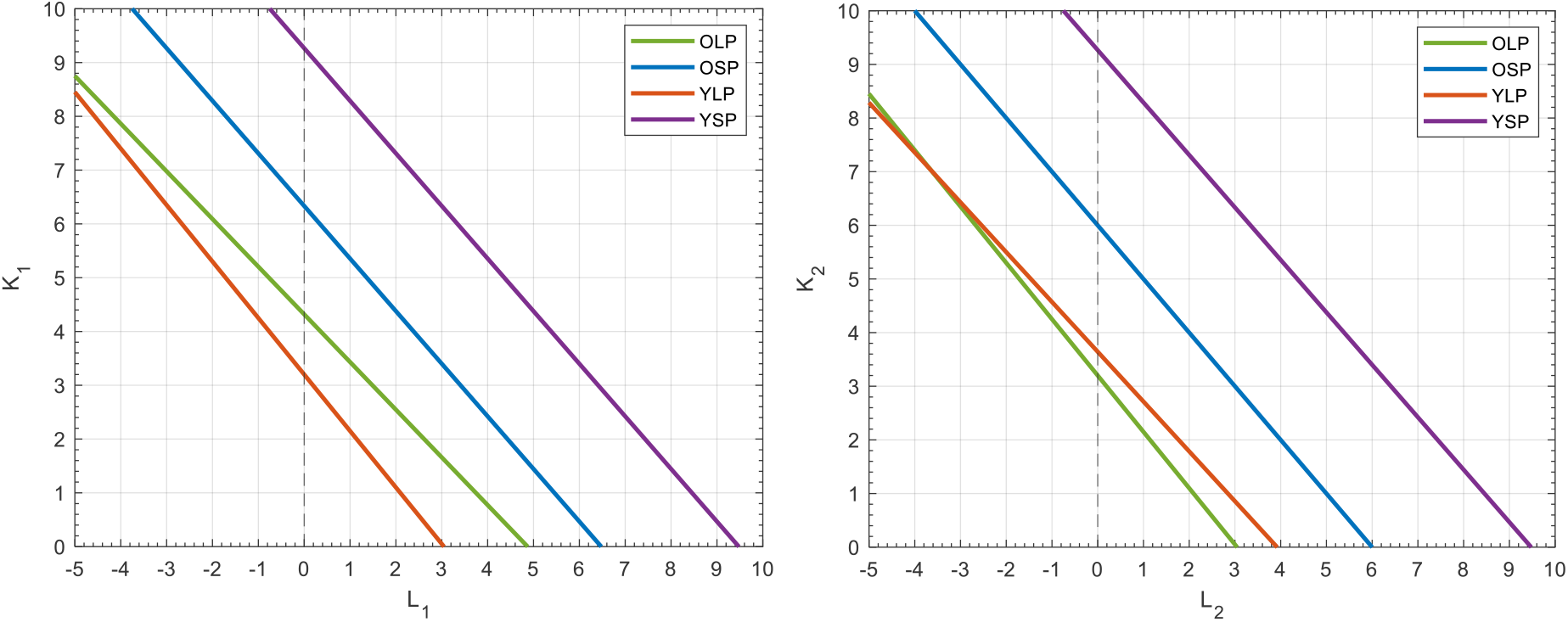
Coupling strength within and between neuronal subpopulations of the SCN. (A) The relation between the coupling strength (*K*_*1*_) within the medial oriented cluster and the interaction strength (*L*_*1*_) from the lateral oriented cluster to the medial oriented cluster are shown for the different experimental conditions. The green line are old mice in LP, the blue line old mice in SP, the orange line are young mice in LP and the purple line are young mice in SP. There is a range of values for *K*_*1*_ and *L*_*1*_ that result in the same synchronization as observed in the bioluminescence data. (B) The same as Fig A for the coupling strength (*K*_*2*_) within the lateral oriented cluster and the interaction strength (*L*_*2*_) from the medial oriented cluster to the lateral oriented cluster

Taken more general we can describe the relation between *K*_*1*_ and *L*_*1*_ as the linear line

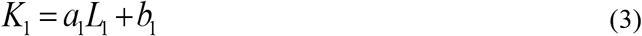

in which 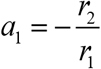 and *b*_*1*_ is only dependent on *r*_*1*_ in an exponential manner. Therefore, when *r*_*1*_ is greater than *r*_*2*_ the slope of the line is greater than -1 and when *r*_*1*_ is smaller than *r*_*2*_ the slope of the line is smaller than -1. Furthermore, when *r*_*1*_ increases the line shifts vertically upwards and when *r*_*1*_ decreases the line shifts vertically downwards. The relationship between *K*_*2*_ and *L*_*2*_ can be described in the same way, by interchanging the role of *r*_*1*_ and *r*_*2*_.

From our available experimental data it is difficult to obtain precise values for *K*_*1*_, *K*_*2*_, *L*_*1*_ and *L*_*2*_. We found that the synchronization in the neuronal subpopulations is always higher than the synchronization of the SCN as a whole and from the relation between *r* and *K* we know that *K* goes to infinity when *r* reaches 1. This suggests that the coupling strength within the neuronal subpopulations is higher than the coupling strength in the SCN in general. However, we were not able to measure the single cell traces from the neuronal subpopulations independent of each other. Therefore it is unclear whether the increased level of synchronization within the neuronal subpopulations is caused by an increase in coupling strength within the clusters or due to the interaction strength between the clusters or due to a combination of both.

Although we cannot estimate the precise values for *K*_*1*_, *K*_*2*_, *L*_*1*_ and *L*_*2*_ for the different experimental conditions, we can already learn more about the coupling strength in the SCN from the relations between *K* and *L* for the different experimental conditions. For instance, for the complete range of *L*, the lines from young mice in SP and LP are further apart than the lines from old mice in SP and LP. This indicates that the range over which young mice can adapt their coupling strength is larger than the range over which old animals can adapt their coupling strength.

## Discussion

In this study we analyzed single-cell PER2::LUC gene expression rhythms of SCN neurons to determine the synchronization levels in the SCN as a whole and within the neuronal subpopulations in the SCN for young and old mice in long and short photoperiod. By use of the Kuramoto model we identified that the SCN of old animals is less able to adjust to a short photoperiod because of an inability to respond to short photoperiod with an increase in coupling strength. There is no difference between young and old animals in long photoperiod, when only a low degree of coupling is required. Hence, exposure to short photoperiod is not a successful strategy in order to boost the rhythm of old animals.

The extended Kuramoto model appeared to be useful to determine the coupling strengths between neurons in the SCN based on PER2::LUC data, once the noise component was separated from the coupling strength. From the relation between *K* and *L* we have found, we can make two statements regarding coupling strength in the SCN. First, for the whole range of interaction strengths (*L*_*1*_ and *L*_*2*_), the coupling strength (*K*_*1*_ and *K*_*2*_) is higher in SP than LP in both young and old mice. Higher coupling strength in SP than LP confirms that the higher synchronization seen in SP is supported by changes in coupling strength. Second, the range over which young mice can adapt their coupling strength between SP and LP is larger for both subpopulations than the range over which old mice can adapt their coupling strength between SP and LP. This indicates that old mice are less capable of adapting to different photoperiods. This is in agreement with previous data [15], showing that old mice had behaviorally a strongly reduced ability to adapt to different photoperiods.

There will always be variability between animals within an experimental group. For neuronal synchronization in the SCN it is known that there is experimentally more variability in the level of phase coherence between old mice than between young mice as well as there is more variability between mice in LP than in SP [15,25]. This is in agreement with our results for neuronal synchronization in the SCN. In old mice there is a larger variability in synchronization within any experimental condition than in young mice and for both young and old mice there is a larger variability in synchronization in LP than in SP.

Previous studies showed that synchronization between neurons was increased in short photoperiod and decreased in long photoperiod [7,10]. In Fig 2A can be seen that for young mice the upper bound of the coupling strength in LP is lower than the lower bound of the coupling strength in SP. Thus, there is no overlap in the coupling strength levels between SP and LP, meaning that the coupling strength in SP is always higher than in LP. For old animals the ranges of the coupling strength do overlap between SP and LP. However, the lower bound of the coupling strength in SP lies within the upper half of the range for the coupling strength in LP, indicating that it is plausible that the coupling strength is higher in SP than LP, also for old animals. The noise is approximately the same in all experimental conditions, which is to be expected since the thermal environment of the neurons does not change between experimental conditions. This finding indicates that the increment in variability in old mice do not result from an increment in noise level.

The order parameter, representing the synchronization, was normalized to obtain a value between 0 and 1, in which 0 means that the phases of the single-cells are randomly distributed and 1 implies perfect synchrony [26,27]. A limitation of the extended Kuramoto model is that coupling strength would become infinite when the neuronal synchronization of the SCN is 100%. This problem is theoretical rather than practical: due to the differences in intrinsic characteristics of the neurons and noise in the system, perfect synchronization will never be reached [28].

One unique property of the extended Kuramoto model used in this study is that the coupling strengths between and within the two communities can vary from each other. From chemical coupling it is known that the dorsal SCN receives strong input from the ventral SCN, whereas the ventral SCN receives scarce input from the dorsal SCN [29,30]. This can be taken into account when using the model by taking different values for *L*_*1*_ and *L*_*2*_ to make the simulations more realistic. However, we do not know whether the constraints for chemical and molecular coupling are the same. Identifying constraints for coupling strength between (and within) communities could help in further specifying the dynamics in the SCN, using a modeling approach.

Previous modeling work by Myung and Pauls [31] describes the interaction between two functional oscillators: one in the dorsal and one in the ventral SCN. Their work pioneered in showing the existence of repulsive coupling from the ventral part of the SCN to the dorsal part of the SCN and attractive coupling from the dorsal part of the SCN to the ventral part of the SCN. Myung and Pauls furthermore suggested that the repulsive coupling strength is higher in LP than in SP, creating a wider peak time dispersion between neurons in LP. We could translate the results of Myung and Pauls as constraints into our model, but then we have to keep in mind that the model of Myung and Pauls was not aimed nor designed to describe coupling among the neuronal oscillators within the dorsal or within the ventral SCN, but was aimed to describe coupling only between the dorsal and ventral SCN. The addition of parameters for the coupling strength within neuronal subpopulations makes our model more realistic, but makes it computationally more complicated. Their ventral cluster would approximately match with our medially oriented cluster and their dorsal cluster would approximately match with our laterally oriented cluster. Increasing the repulsive interaction strength between the medially oriented cluster and the laterally oriented cluster in LP compared to SP is possible in our model. This would have as result that the differences in coupling strength within the clusters (*K*) between photoperiods would decrease in comparison with a situation where the coupling between clusters (*L*) would be similar between photoperiods (Fig 6).

**Fig 6.**
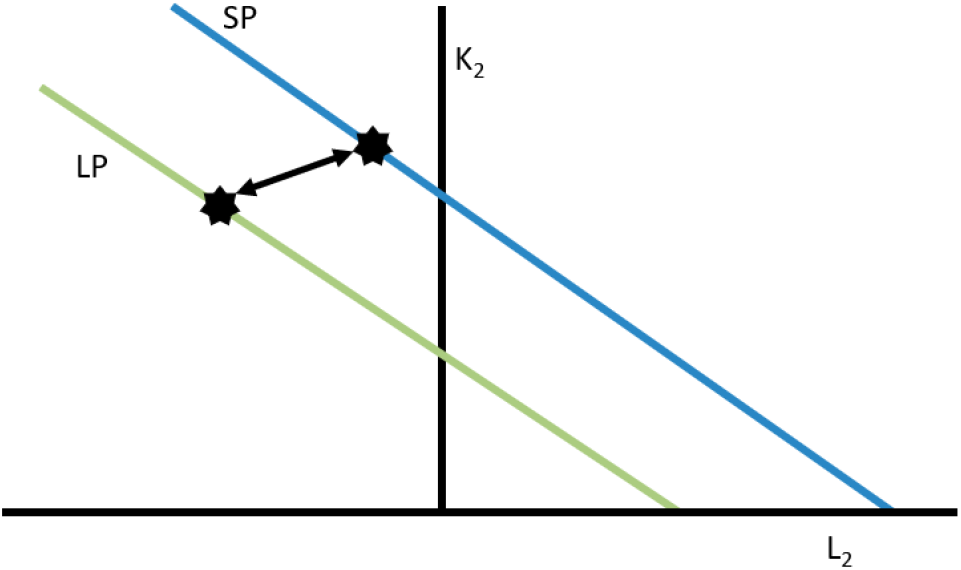
Constrain for the Kuramoto model. A study by Myung and Pauls [31] suggested that the negative coupling strength from the ventral part to the dorsal part of the SCN is stronger in LP than in SP. The black stars show how this would influence our model. The difference in coupling strength (K_2_) would be small between LP (green line) and SP (blue line).

To recapitulate, with the extended Kuramoto model we could determine the coupling strengths between neurons in the SCN, after we measured the synchronization of the neurons, if there was a constant thermal environment (which can be provided). We found evidence that coupling strength within and between subpopulations contributes to photoperiod induced changes in the phase relationship between neurons. In long photoperiod we found lower coupling strengths, and in short photoperiod higher coupling strengths both between and within populations. In young mice, the coupling strengths are higher during short photoperiod than in old mice, as aged mice appear to have a reduced capacity to reach a higher coupling strength in the SCN. The extended Kuramoto model appeared to be highly suitable to determine network properties of the SCN, that are not directly measurable, but can be derived on the basis of available empirical data.

## Methods

### Bioluminescence Imaging and Analysis

To obtain the parameters for the Kuramoto model, the PERIOD2::LUCIFERASE (PER2::LUC) gene expression data from the studies [7,15] was used. The dataset consisted of bioluminescence data from young (4-8 months) and old (22-28 months) homozygous PER2::LUC mice entrained to either long photoperiod (LD 16:8) or short photoperiod (LD 8:16). For details on the data collection see Buijink et al. In short, mice were killed 1 to 3 h before lights-off. The brain was dissected and the SCN was sliced in coronal slices with a VT 1000S vibrating microtome (Leica Microsystems, Wetzlar, Germany). Slices containing the SCN were optically identified and placed in a petri dish. The dish was transferred to a temperature-controlled (37°C) light-tight chamber, equipped with an upright microscope and a cooled charge-coupled device camera (ORCA-UU-BT-1024, Hamamatsu Photonics Europe, Herrsching am Ammersee, Germany). Bioluminescence images were collected with a 1-h time resolution.

To analyze the time series of bioluminescence images a custom-made MATLAB-based (Mathworks, Natick, MA, USA) program was used, as described in [7]. Briefly, groups of adjacent pixels with luminescence intensity above the noise level were defined as regions of interest (ROIs). Each ROI is referred to as a ‘single cell’. The average bioluminescence of all pixels in each ROI was calculated for the image series, which resulted in the bioluminescence traces representing PER2::LUC expression for all single-cell ROIs. For the analysis of rhythm characteristics, such as peak time and period, the raw PER2::LUC expression traces were smoothed and resampled to one data point per minute. Only single-cell traces containing at least three cycles with a period length between 20-28 hours were included for further analysis.

The phase distribution and the Kuramoto order parameter (*r*) were calculated for all SCN slices. Phase distribution was defined as the standard deviation (*SD*) of the peak times from all cells in a slice of the specified cycle in vitro. The order parameter is a measure for synchronization and is based on the relative phase of the single cells. The order parameter was determined by first calculating the mean peak time 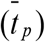 of PER2::LUC expression of all cells (*j* = 1 … *N*) for the specified cycle:

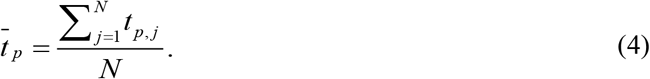

Then the relative phase of each cell was approximated by first subtracting the peak time of the individual cell from the averaged peak time of all cells to get the relative peak time and then converting the relative peak time to its relative phase (*θ*_*r*_) :

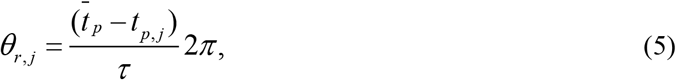

where *τ* is the period in hours. The relative phase can be approximated because the sin(*x*) function is linear for small *x* and the relative peak times are small in comparison with the period. Thereafter, the relative phase was transformed with Euler’s formula and the absolute value was taken to get the order parameter *(r)*:

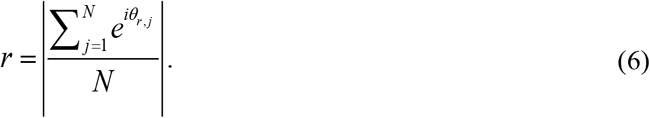

The order parameter can take values between 0 and 1, in which 0 means that the neurons are completely unsynchronized and 1 means perfect synchrony.

### Community Detection

To identify functional clusters in the SCN neuronal network, we used a community detection method that was previously described by [21]. In brief, from the raw time series of PER2::LUC bioluminescence traces a cross-correlation matrix was constructed. Next, with the use of random matrix theory, the global (SCN-wide) and local (neuron-specific) noise components were filtered out of the cross-correlation matrix. Clusters were detected with optimally contrasted functional signature, resulting in a positive overall correlation within clusters and a negative overall correlation between clusters, relative to the global SCN activity. Although the clustering algorithm was not bound to a pre-defined number of groups, the community detection method results consistently in two main groups of cells with a robust spatial distribution. The spatial distribution differed slightly for the anterior and posterior slices [7,15]. Hence, the resulting clusters were visually labeled as ventromedial and dorsolateral in the anterior SCN and as medial and lateral in the posterior SCN slices.

### Kuramoto model

To model the SCN we used a Kuramoto model. The Kuramoto model is a simple model that only contains phase information [22]. First we used a one-community Kuramoto model to find an upper and lower bound for the coupling strength in the different experimental conditions. Furthermore we used the one-community Kuramoto model to estimate the amount of noise in the model. The noise term represents the thermal environment of the SCN (i.e., external noise), which should be the same in all experimental conditions. With use of the one-community model we show that the amount of noise is indeed approximately the same in the different experimental conditions. The same amount of noise between the different experimental conditions was a requirement to extend to a two-community Kuramoto model, such that the influence of the noise could be separated from the influence of the coupling strength. We used the two-community Kuramoto model to find the relationship between the coupling strength within each subgroup and the coupling strength between the two subgroups. Here we took the upper and lower bounds for the coupling strength as found with the one-community model into consideration.

### One-community Kuramoto model

In the one-community Kuramoto model we consider one-community of *N* oscillators. Each oscillator corresponds to a neuron in the SCN. The oscillators interact with a strength *K* which gives a mean-field interaction strength *K/N*. The phase angles of the oscillators are denoted by *θ*_*i*_, *i=1*, …, *N* and represent the state of the neuron. To simplify the model we have set the natural frequency of all oscillators to zero. Since any constant frequency can be rotated out by changing the frame of reference of the system, any constant average natural frequency can be chosen [23]. The equation for a single representative neuron is given by:

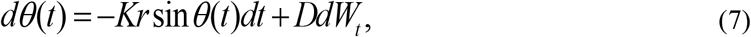

Where *D* is the noise strength and *W*_*t*_ is a standard Brownian motion. The noise can be understood as the effect of the thermal environment of the SCN or as time-dependent variations in the natural frequencies of individual oscillators. Now we will integrate the SDE from 0 to *T* giving:

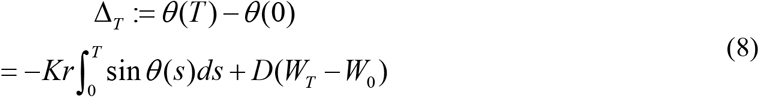

Which, when taking the expectation leads to

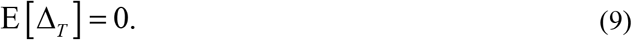

From Itô calculus we can calculate

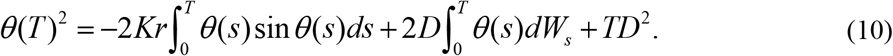

In order to find upper and lower bounds on the noise strength we will take the expectation of Δ_*T*_ using two expansions of the sinusoidal function and we will take the initial phase to be zero so that *θ* (0) = 0. Taking 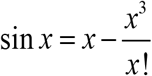 gives

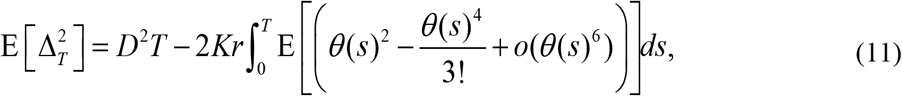

Which implies that

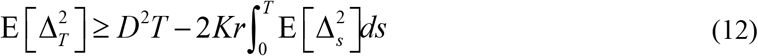

And since we are in stationarity this gives an upper bound for the noise strength *(D*_+_)

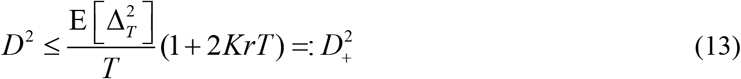

Using one more term in the expansion for sin gives

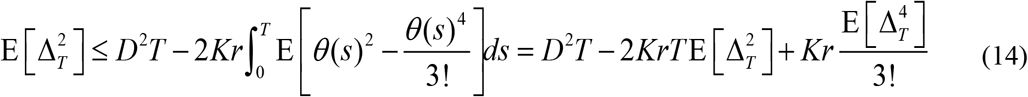

so that the noise strength is bounded from below *(D*_-_) by

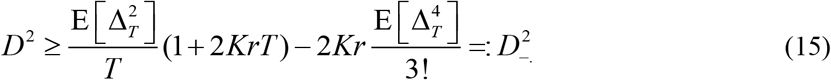

Since we have observations of Δ_*T*_ we are able to numerically calculate upper and lower bounds for the noise strength in terms of the interaction strength *K*. This holds in the case that sine is approximated well by the expansion used, which we posit to be the case since the spread of the phases around the average is small relative to the size of the entire cycle.

In order to do this we need unbiased estimators of the second and fourth moments. Since the mean is zero, the fourth moment is equal to the fourth central moment for which an unbiased estimator is given by the fourth h-statistic

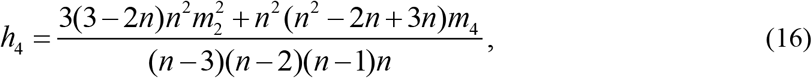

where *n* is the sample size and *m*_*p*_ is the *p*^*th*^ sample central moment given by

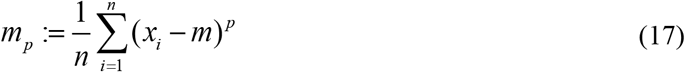

With *m* the sample mean. An unbiased estimator for the variance is

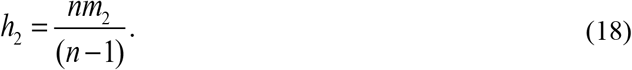

Now if we want to calculate the parameter for a single community we must solve the equation

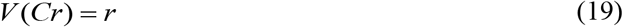

where

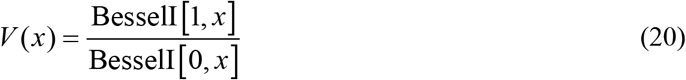

and Besse1I[0, *x*] and Besse1I[1, *x*] are modified Bessel functions of the first kind. From the bioluminescence data we have calculated *r* so that we can use numerical methods (like FindRoot in MATHEMATICA) to solve for *C*. In the one-community model 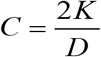 so that we can find upper and lower bounds for *K*

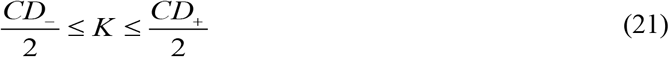

Now both *D*_-_ and *D*_+_ depend on *K* so that we find

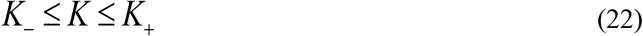

with

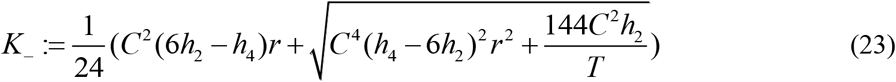

and

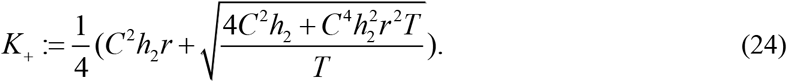

### Two-community Kuramoto model

The one-community Kuramoto model was elaborated to a two-community model in which both communities consist of *N* oscillators. The oscillators in the same community interact with strength *K* and oscillators in different communities interact with strength *L*. The phase angles of the oscillators in the first community are denoted by *θ*_*1,i*_, *i=1*, …, *N* and in the second community by *θ*_*2,j*_, *j=1*, …, *N*. The equations governing their evolution are then:

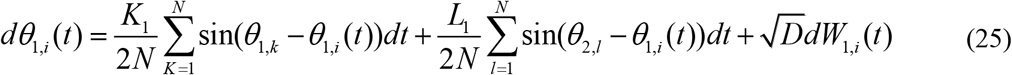

and

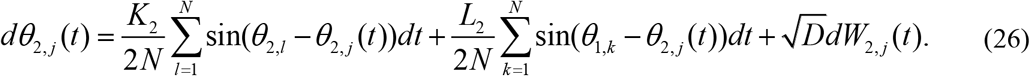

From the one-community Kuramoto model we found that *D* does not depend on the synchronization levels and that *D* is close to 1 for all experimental conditions. Therefore we take *D* as a constant in the two-community Kuramoto model. Furthermore we made the assumption that the average phase is the same in both communities (i.e. *ψ*_*1*_ = *ψ*_*2*_ = *0*). Now we can calculate the relationship between *K*_*1*_ and *L*_*1*_ and between *K*_*2*_ and *L*_*2*_ by solving the equations

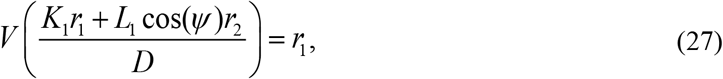

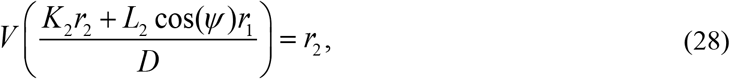

in the same way we did for the one-community Kuramoto model. In the above equations *K*_*1*_ and *K*_*2*_ represent the coupling strengths within respectively subpopulations 1 and 2. *L*_*1*_ and *L*_*2*_ represent the interaction strength between subpopulations, where *L*_*1*_ is the strength from subpopulation 2 to subpopulation 1 and *L*_*2*_ is the strength from subpopulation 1 to subpopulation 2. *r*_*1*_ and *r*_*2*_ are the order parameters in respectively subpopulation 1 and 2 and *ψ* is the phase difference between the subpopulations.

## Acknowledgements

We thank Conrado da Costa for helping with the calculations for the upper and lower bounds for K and D and we thank Pablo Villegas for valuable comments on an earlier version of the article.

## Author Contributions

**Conceptualization:** JMM, JHM and JHTR

**Data curation:** AWB and JHTR

**Formal analysis:** AWB, JMM and SA

**Methodology:** AWB, JMM, SA and JHTR

**Writing – original draft:** AWB and JHTR

**Writing – review & editing:** AWB, JMM, SA, JHM and JHTR

## Supporting Information

**Fig S1.**
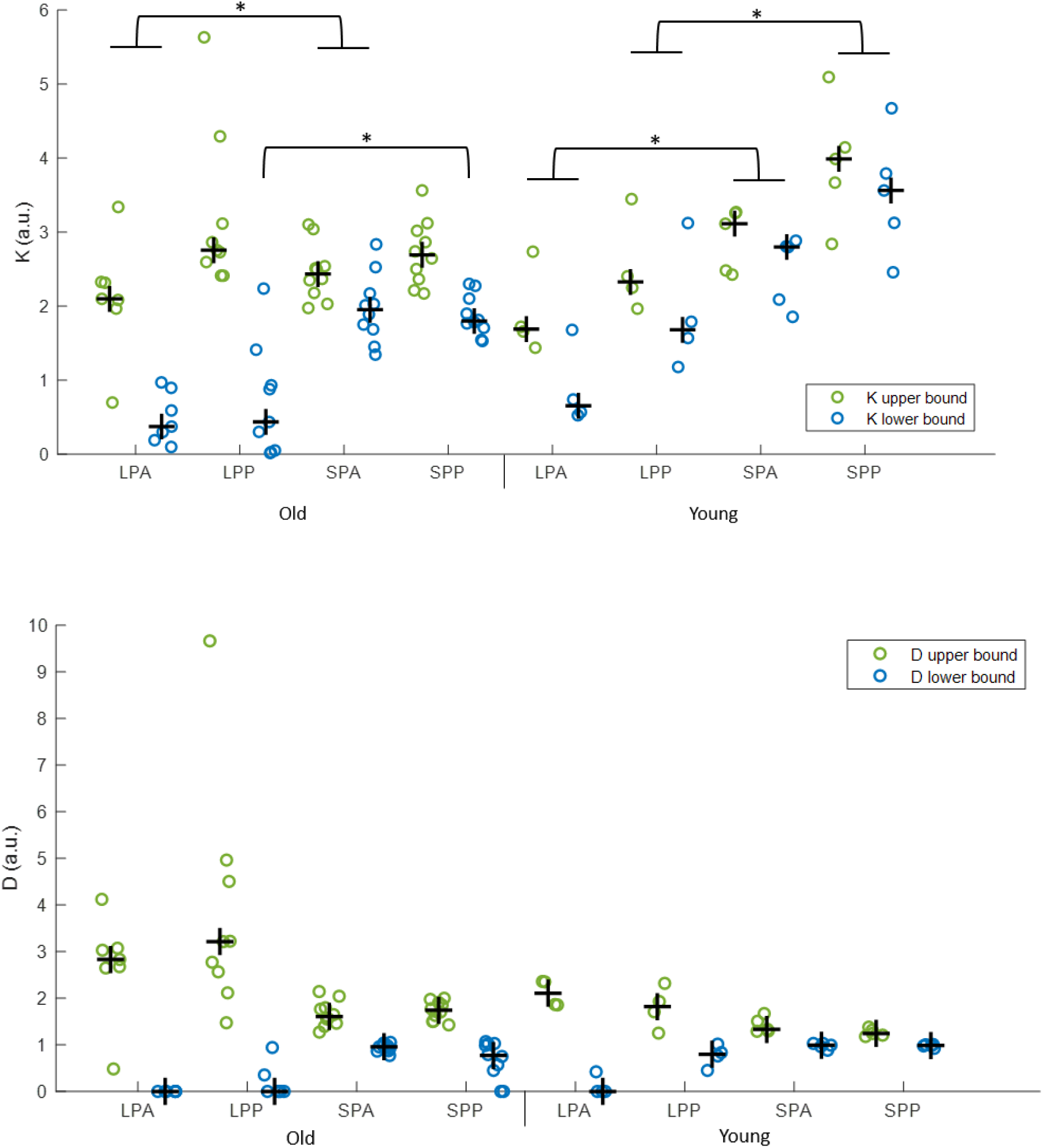
Estimation of the upper and lower bound of K and D in different experimental conditions. Upper (blue dots) and lower (green dots) bound of K (top plot) and D (bottom plot) estimated with the Kuramoto model for anterior (A) and posterior (P) slices in short (SP) and long photoperiod (LP) in old and young mice. The black cross indicates the median; *p<0.05

